# *DriverGroup*: A novel method for identifying driver gene groups

**DOI:** 10.1101/2020.04.23.058719

**Authors:** Vu VH Pham, Lin Liu, Cameron P Bracken, Gregory J Goodall, Jiuyong Li, Thuc D Le

## Abstract

**Motivation:** Identifying cancer driver genes is a key task in cancer informatics. Most exisiting methods are focused on individual cancer drivers which regulate biological processes leading to cancer. However, the effect of a single gene may not be sufficient to drive cancer progression. Here, we hypothesise that there are driver gene groups that work in concert to regulate cancer and we develop a novel computational method to detect those driver gene groups.

**Results:** We develop a novel method named *DriverGroup* to detect driver gene groups by using gene expression and gene interaction data. The proposed method has three stages: (1) Constructing the gene network, (2) Discovering critical nodes of the constructed network, and (3) Identifying driver gene groups based on the discovered critical nodes. Before evaluating the performance of *DriverGroup* in detecting cancer driver groups, we firstly assess its performance in detecting the influence of gene groups, a key step of *DriverGroup*. The application of *DriverGroup* to DREAM4 data demonstrates that it is more effective than other methods in detecting the regulation of gene groups. We then apply *DriverGroup* to the BRCA dataset to identify coding and non-coding driver groups for breast cancer. The identified driver groups are promising as several group members are confirmed to be related to cancer in literature. We further use the predicted driver groups in survival analysis and the results show that the survival curves of patient subpopulations classified using the predicted driver groups are significantly differentiated, indicating the usefulness of *DriverGroup*.

**Availability and implementation:** *DriverGroup* is available at https://github.com/pvvhoang/DriverGroup

**Supplementary information:** Supplementary data are available at *Bioinformatics* online.

## 1 Introduction

It is important to identify cancer drivers and their regulatory mechanisms since cancer drivers play a critical role in the initialisation and progression of cancer. Understanding cancer drivers is beneficial for the design of effective cancer treatments too. Thus, several computational methods have been developed to discover cancer drivers (Gonzalez-Perez and Lopez-Bigas, 2012; Tamborero *et al.*, 2013; Reimand and Bader, 2013). In general, we can catergorise the existing methods into mutation-based methods and network-based methods. Mutation-based methods detect cancer drivers based on mutations and their characteristics, such as functional impact used by OncodriveFM (Gonzalez-Perez and Lopez-Bigas, 2012), recurrence by OncodriveCLUST (Tamborero *et al.*, 2013), enrichment in externally defined regions by ActiveDriver (Reimand and Bader, 2013) and mutual exclusivity by CoMEt (Leiserson *et al.*, 2015). Network-based methods estimate the role of genes in a biological network and may combine the results with additional information such as mutations to discover cancer drivers. The representatives for the network-based approach are DawnRank (Hou and Ma, 2014), DriverNet (Bashashati *et al.*, 2012), CBNA (Pham *et al.*, 2019), and NBS (Hofree *et al.*, 2013).

All of the above methods only identify single genes as cancer drivers. However, the effect of a single gene may not be sufficient to regulate cancer. In this paper, we introduce the concept of “driver gene group”, which is a set of genes that work in concert to regulate cancer or cancer markers. There is evidence showing that genes work together to regulate the same targets and the regulation of individual genes might not have significant impacts (Cursons *et al.*, 2018; Karim *et al.*, 2016). Furthermore, researchers have started to conduct wet-lab experiments to investigate the regulation by groups of genes in biological processes (Cursons *et al.*, 2018). All of these highlight the importance of studying biological components working in groups, including groups of cancer drivers which collaboratively regulate targets to drive cancer.

The driver gene groups are different from the gene modules which are studied by the recent methods such as WeSME (Kim *et al.*, 2017), MEMo (Ciriello *et al.*, 2012), and iMCMC (Zhang *et al.*, 2013). WeSME (Kim *et al.*, 2017) discovers cancer drivers by using statistical tests to evaluate the mutual exclusivity of mutations of gene pairs and the pairs whose mutations have a significant mutual exclusivity are considered as modular candidate drivers. Similar to WeSME, MEMo (Ciriello *et al.*, 2012) also uses mutual exclusivity of gene mutations in detecting cancer drivers. However, instead of testing the mutual exclusivity of mutations of gene pairs, MEMo tests the mutual exclusivity of mutations of genes in subnetworks. The subnetworks include genes which are recurrently altered in samples and likely to belong to the same pathway, and the subnetworks whose mutations have a significant mutual exclusivity are considered as modular candidate drivers. iMCMC (Zhang *et al.*, 2013) uses somatic mutations, copy number variations, and gene expression to build a gene network. It then extracts modules (i.e. coherent subnetworks with large weights in both edges and nodes) from the network. The driver modules are the ones having significant results in the random test and the exclusivity test. The random test is for evaluating the significance of results and the exclusivity test is for assessing if a module exhibits the pattern of mutually exclusive genomic alterations.

Although the above methods detect modules of cancer drivers, the mutation in just one member of a module is sufficient to trigger cancer development and the members of each module may not work jointly to regulate targets to drive cancer (Kim *et al.*, 2017). However, the idea of driver gene groups is that all genes in a group collaboratively drive cancer. In addition, these methods only deal with coding genes while cancer drivers may be non-coding genes since a large portion of mutations may exist in non-coding regions (Yang *et al.*, 2016a), and non-coding genes can regulate gene targets to drive cancer (Puente *et al.*, 2015; Weinhold *et al.*, 2014). Thus, there is a strong need for novel methods to identify both coding and non-coding driver gene groups of which the members of each driver group work in concert to progress cancer.

In this paper, we propose a novel method named *DriverGroup* to identify both coding and non-coding driver gene groups. Due to the fact that proliferation is associated with cancer development (Lopez-Saez *et al.*, 1998; Feitelson *et al.*, 2015) and proliferation genes are related to the prognosis of cancer patients (Li *et al.*, 2018), we identify driver gene groups by detecting groups of genes which collaboratively regulate proliferation genes (i.e. groups of genes which regulate more target genes than the union of the target genes of individuals in the groups).

We base on the gene network and its critical nodes (i.e. nodes playing a central role in controlling the whole network) to identify driver gene groups. Because of the important role of critical nodes, we consider them as members of driver gene groups. Inspired by the Influence Maximisation (IM) problem (Gong *et al.*, 2016; Yang *et al.*, 2016b) which identifies k-seed sets (i.e. sets have k seed nodes) with the maximum influence in a network, we develop novel algorithms to compute the influence of a group of critical nodes on the proliferation genes. At the end, a driver gene group is a maximal subset of critical nodes which have the maximum impact on the proliferation genes, i.e. adding or removing one critical node from the subset will decrease the impact of the subset.

Before evaluating *DriverGroup* in identifying driver gene groups, we firstly assess its ability in discovering the influence of the gene groups in a network. We use the DREAM4 data obtained from the DREAM4 In Silico Network Challenge (Marbach *et al.*, 2010; Schaffter *et al.*, 2011). We compare *DriverGroup* with jointIDA (Nandy *et al.*, 2017), a method used to estimate the joint effects of a group of variables on other variables by knocking down all regulators at the same time, and the random method. As a result, our method outperforms both jointIDA (Nandy *et al.*, 2017) and the random method in most cases. We use the BRCA dataset for identifying driver gene groups and several members of the driver groups predicted by *DriverGroup* are confirmed to be related to cancer by literature, suggesting the biological meaning of the findings of the proposed method. We go further to analyse the ability of the driver groups predicted by *DriverGroup* in prognosis. The results show that the subtypes identified based on the predicted driver groups have significant prognostic values for survival analysis (i.e. p-values less than 0.05), indicating that the driver groups identified by *DriverGroup* may have important clinical implications for cancer treatment. We also apply *DriverGroup* to the study of synthetic lethality and miRNA driver groups of epithelial-mesenchymal transition (EMT). Several predicted synthetic lethality gene pairs are in the existing database and many members of the predicted miRNA driver groups for EMT are in the list of EMT miRNAs (i.e. miRNAs related to EMT). These results demonstrate that *DriverGroup* is useful not only in exploring driver gene groups but also in predicting driver groups for other processes such as EMT. The results also show the potential of *DriverGroup* as a framework for studying molecular mechanisms of the progression of cancer and other diseases.

## 2 Datasets and method

### 2.1 Datasets

In this study, we use the BRCA dataset of TCGA (The Cancer Genome Atlas Research *et al.*, 2013). This dataset contains the expression data of miRNAs, TFs, and mRNAs of tumour/normal samples. The tumour samples are used to identify edges of the gene network and the normal samples are used to compute node weights. The TF list, which is used to detect which genes are TF genes in the expression dataset, is obtained from Lizio *et al.* (2017). We also use interaction data (i.e. target binding information), including PPIs (Vinayagam *et al.*, 2011), miRNA-TF/mRNA interactions (miRTarBase 6.1 (Chou *et al.*, 2016), TarBase 7.0 (Vlachos *et al.*, 2015), miRWalk 2.0 (Dweep and Gretz, 2015), and TargetScan 7.0 (Agarwal *et al.*, 2015)), and TF-miRNA interactions (TransmiR 2.0 (Wang *et al.*, 2010)) to refine the built gene network. In addition, to evaluate the performance of *DriverGroup* in detecting the influence of gene groups in a network, we use DREAM4 data obtained from the DREAM4 In Silico Network Challenge (Marbach *et al.*, 2010; Schaffter *et al.*, 2011). We also use the SynLethDB synthetic lethality database (Guo *et al.*, 2016) for identifying synthetic lethality, the EMT signatures (Tan *et al.*, 2014) and the EMT miRNAs (Cursons *et al.*, 2018) for discovering EMT driver groups. More details of these datasets will be introduced in the following sections. All these datasets are available at https://github.com/pvvhoang/DriverGroup.

### 2.2 DriverGroup

#### 2.2.1 Overview

An overview of *DriverGroup*, the proposed method for identifying driver gene groups, is shown in Fig. 1. *DriverGroup* includes three stages: (1) Constructing the miRNA-TF-mRNA network, (2) Discovering critical nodes in the constructed network, and (3) Identifying driver gene groups. Particulartly, we firstly construct the network using the matched expression data of mRNAs, Transcription Factors (TFs), and miRNAs of a given cohort of cancer patients. Then the directed PPI network (Vinayagam *et al.*, 2011) and the target binding information are used to refine the network by removing those interactions not supported by these databases. Next, we discover critical nodes of the network by applying control theory (Kalman, 1963) and the Network Control method (Liu *et al.*, 2011). The critical nodes play a central role in controlling the whole network. Finally, based on the network and its critical nodes, we identify driver gene groups. The detail of *DriverGroup* is described in the following sections.

**Fig. 1.**
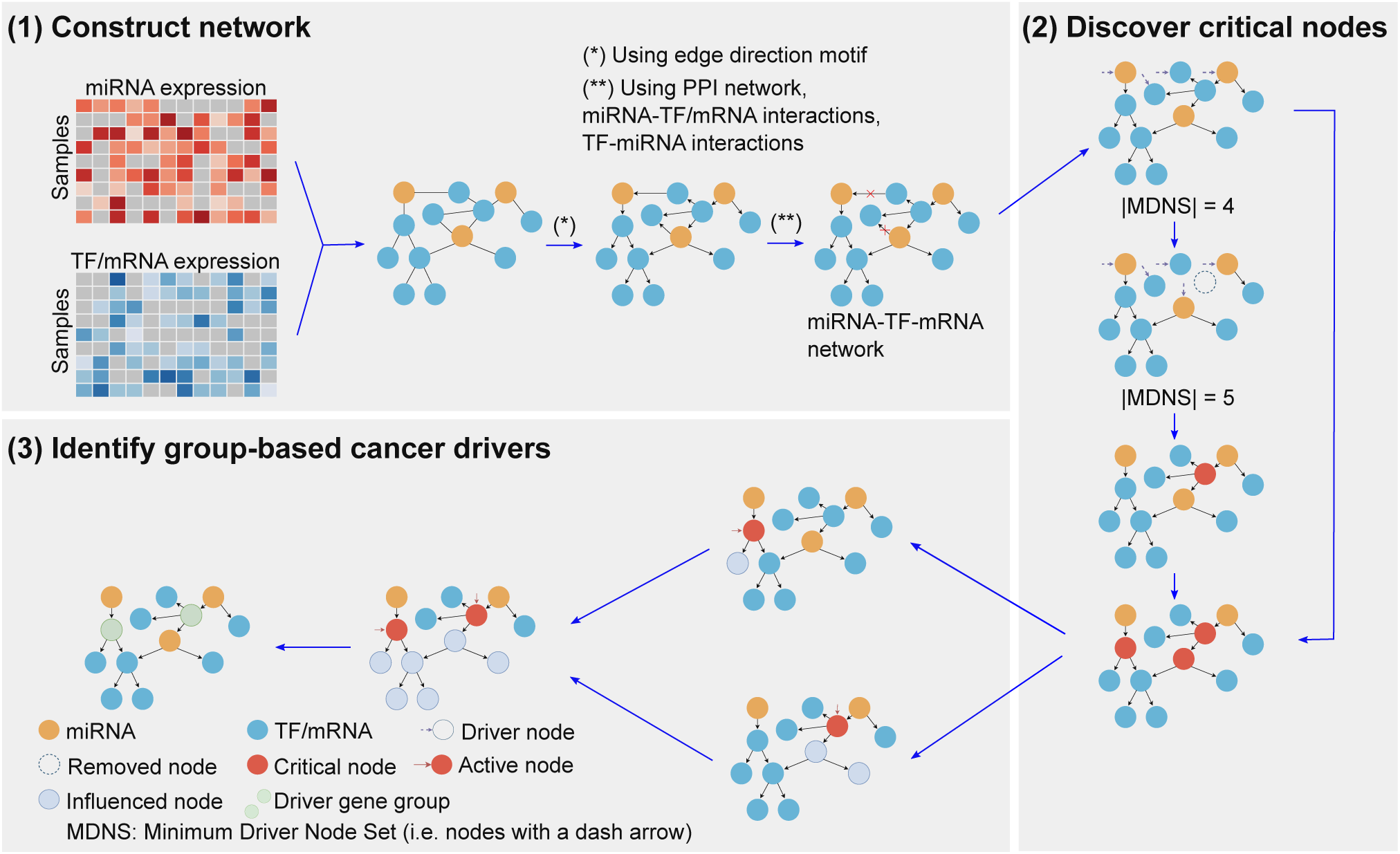
An illustration of *DriverGroup*. (1) Build the network by combining the gene network built from the expression data with the protein-protein interaction (PPI) network and other existing databases, including miRTarBase 6.1, TarBase 7.0, miRWalk 2.0, TargetScan 7.0, and TransmiR 2.0, (2) Discover critical nodes by evaluating the size of Minimum Driver Node Set (MDNS) (i.e. the set of nodes whose size is minimum and it can control the whole network) when a node is removed, and (3) Identify driver gene groups by detecting the influence of groups of critical nodes on the proliferation genes.

#### 2.2.2 Procedure for identifying driver gene groups

##### Stage (1) Constructing the miRNA-TF-mRNA network

To identify driver gene groups, we detect groups of miRNAs and coding genes which jointly impact on the proliferation genes in a gene regulatory network. Since we evaluate both miRNA cancer driver groups and coding cancer driver groups, we construct the network which includes both miRNAs and coding genes (i.e. TFs and mRNAs). It is called the miRNA-TF-mRNA network in this paper. In the first stage, we build the miRNA-TF-mRNA network through the three following steps.

- Step 1a: Prepare the miRNA/TF/mRNA expression data. We obtain the miRNA/TF/mRNA expression data of matched samples from the BRCA dataset (The Cancer Genome Atlas Research *et al.*, 2013). For coding genes, we select genes which are in the PPI network (Vinayagam *et al.*, 2011) or in the proliferation gene list. We select the PPI network as it contains a large amount of cancer driver genes and has been used to identify driver genes by Vinayagam *et al.* (2016). The proliferation genes are retrieved from the biological process of cell population proliferation (GO:0008283). We then use the TF list to detect which genes are TFs in the selected genes. As a result, we have 5,273 mRNAs and 850 TFs. For miRNAs, we select all 1,719 miRNAs from the BRCA dataset. Finally, we extract the expression data of these 1,719 miRNAs, 850 TFs, and 5,273 mRNAs for 747 tumours and 76 normal samples.
- Step 1b: Build the miRNA-TF-mRNA network. We firstly build a miRNA-TF-mRNA network based on the miRNA/TF/mRNA expression data of tumour samples. A node of the network is a miRNA, a TF, or a mRNA. The node weight is the absolute difference of average expression of that node between tumour and normal states. The node weight indicates the cost to change the state of a node from normal to tumour. The bigger the weight of a node is, the higher the cost required to change between the states. An edge between two nodes is added if the absolute Pearson correlation coefficient between them (calculated based on expression data) is larger than or equal to a threshold, which is the average of the absolute pairwise Pearson correlation coefficients of all node pairs. The edge weight is the absolute value of the correlation coefficient between the two nodes. Edge directions are determined according to the motif shown in Fig. 2. Particularly, miRNAs can regulate TFs/mRNAs, TFs can regulate miRNAs/mRNAs, and TFs/mRNAs can regulate other TFs/mRNAs respectively.
- Step 1c: Refine the gene network. We use the PPIs to refine the expression network. If a TF-TF/mRNA or mRNA-mRNA interaction is in the expression network but not in the PPI network, we remove it from the expression network. We continue to refine the obtained network by using the existing databases. Particularly, miRNA-TF/mRNA interactions are refined with miRTarBase, TarBase, miRWalk, and TargetScan. TF-miRNA interactions are refined with TransmiR. Because the obtained network is based on both the expression data of a particular cancer type and the existing databases, it is more reliable and specific to that cancer type. The final network includes 7,842 nodes (1,719 miRNAs, 850 TFs, and 5,273 mRNAs) and 171,459 edges (23,037 miRNA-TF, 105,019 miRNA-mRNA, 30,096 TF-miRNA, 1,235 TF-TF, 815 TF-mRNA, and 11,257 mRNA-mRNA).

**Fig. 2.**
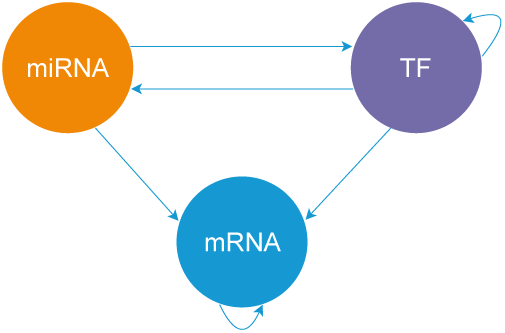
Motif of the edge directions of the miRNA-TF-mRNA regulatory network. In the miRNA-TF-mRNA regulatory network, miRNAs can regulate TFs/mRNAs, TFs can regulate miRNAs/mRNAs, TFs/mRNAs can regulate other TFs/mRNAs respectively.

##### Stage (2) Discovering critical nodes in the network

According to the Network Control method (Liu *et al.*, 2011), any network can be controlled fully by a minimum set of nodes of the network, called a Minimum Driver Node Set (MDNS). The detail of the Network Control method is discussed in Section 2.2.3. Applying this Network Control idea, we discover a MDNS of the miRNA-TF-mRNA network built in Stage (1) above. Based on the discovered MDNS, we then detect critical nodes of the network. A critical node is a node whose absence increases the size of the MDNS. In other words, when a critical node is removed from the network, we need a bigger MDNS to fully control the network. Thus, critical nodes play the central role in the network and we consider them as members of potential driver gene groups. This stage is illustrated in Part (2) in Fig. 1.

##### Stage (3) Identifying driver gene groups

In the last stage, we identify driver groups with the steps below.

- Step 3a: Estimate the influence of groups of critical nodes on the proliferation genes. This step includes the following two substeps.
  1. Form k-way combinations of the selected critical nodes. As we aim to detect nodes which have high influence on the proliferation genes in the network, we focus on nodes with higher out degrees. Out degree of a node is the number of edges going out from that node. We firstly rank critical nodes of the miRNA-TF-mRNA network in descending order of node out degree. We select top n nodes from the ranked list then define k-way combinations of these top n nodes (*k* ∈ {1,…, *n*}).
  2. Evaluate influence of the k-way combinations on the proliferation genes. Influence is indicated by the number of proliferation nodes. Adopting the idea of Influence Maximisation (IM) (Gong *et al.*, 2016; Yang *et al.*, 2016b) for detecting k-seed sets having the maximum impact in a network, we propose a novel algorithm to assess the impact of a group of critical nodes on the proliferation genes. The detail of the proposed algorithm is discussed in Section 2.2.4. Before using the proposed algorithm to evaluate the influence of the k-way combinations, we firstly normalise the node weights and the edge weights of the network so that the weight of a node and the total weight of edges going into a node are in the range from 0 to 1 to make possible to compare the weights in the algorithm. The normalised weight of a node is equal to the original node weight divided by the largest node weight. For edge weights, we firstly find each node’s total incoming edge weight, then find the largest among all these total weights. We normalise an edge weight by dividing it by the largest total weight found. We apply the proposed algorithm to evaluate the influence of each of the k-way combinations of critical nodes. The output of this step is the number of proliferation nodes for each k-way combination (*k* ∈ {1,…, *n*}).
- Step 3b: Identify driver gene groups. In this step, we identify the maximal k-way combinations and regard the identified maximal combinations are the driver gene groups. A k-way combination g (*k* ∈ {1,…, *n*}) is maximal if the (k+1)-way combination obtained by adding to g a critical node has the same or lower influence than g. More details are in Section 2.2.4.

#### 2.2.3 Controllability of complex networks

According to the Network Control method (Liu *et al.*, 2011), any directed network can be controlled by a subset of nodes in the network, known as driver nodes of the network. The method to identify driver nodes is described as follows.

Suppose that we have a directed network with *N* nodes *x*_1_, …, *x*_*N*_. The matrix *A*_*N*×*N*_ which captures the interaction strength between nodes can be represented as:

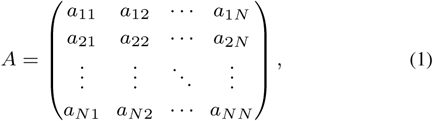

where *a*_*ij*_ indicates the interaction strength of node *j* on node *i* (*i, j* ∈ {1, …, *N*}), and *a*_*ij*_ is 0 if there is not an edge from *j* to *i*.

Let the matrix *B_N×M_* represent the interaction of an external controller on *M* nodes (*M* ≤ *N*) in the network:

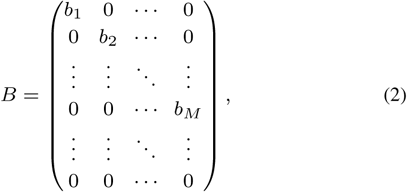

where *b*_*i*_ is the strength of the interaction between the external controller and node *i* (*i* ∈ {1, …, *M*}) in the network.

Let *C*_*N*×*NM*_ be the controllability matrix:

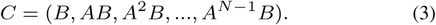

According to Kalman’s controllability condition (Kalman, 1963), the network represented by matrix *A* is controllable through the *M* nodes in matrix *B* if the controllability matrix *C* satisfies the condition:

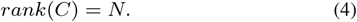

The M nodes are called driver nodes of the network. Intuitively, the rank of the controllability matrix *C* being *N* implies that all *N* nodes of the network can be controlled.

Given the network, we may discover various sets of nodes which satisfy the condition 4, i.e. a network can have multiple sets of driver nodes. In this paper, we focus on the driver node set with the smallest number of driver nodes, called the Minimum Driver Node Set (MDNS). In Stage (2) of *DriverGroup*, we use the condition 4 to detect the MDNS of the miRNATF-mRNA network. We then identify critical nodes of the network by removing one node at a time from the network, and if the MDNS of the network with the node removed is bigger than the MDNS of the original network, the removed node is a critical node.

#### 2.2.4 Influence of groups of nodes

Influence Maximisation (IM) finds a k-seed set that has the maximum influence in a network (Gong *et al*., 2016; Yang *et al*., 2016b). IM is usually used to identify influential users in online social networks (Kempe *et al*., 2003) as described below.

Given a network *G* with *N* nodes and a budget *k*, IM is to find a set *S* containing *k* nodes of *G* (called a k-seed set) which maximises the influence spread over *G*. The influence spread is the number of nodes influenced by a k-seed set and it is denoted as *σ*(*S*). That is:

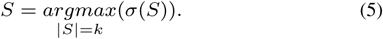

Inspired by IM, we propose the following described method to calculate the influence spread of a k-way combination of critical nodes on the proliferation genes in a miRNA-TF-mRNA network, and then find the maximal combinations as driver groups. In our problem, we do not fix the budget *k* and we evaluate the influence of k-way combinations on the proliferation genes instead of over the whole network.

In general, diffusion models are used to resolve the IM problem (Gong *et al*., 2016). The diffusion models identify the influence spread of a k-seed set over a network by considering that nodes in the k-seed set are active and proposing strategies to activate other nodes in the network. The larger the number of active nodes a k-seed set creates, the more influence it has in the network. These models employ the rules below:

- A node can be active or inactive.
- During the diffusion process, inactive nodes can be activated but active nodes cannot be inactivated.
- The process terminates if no more nodes can be activated.

Independent Cascade (IC) and Linear Threshold (LT) are two popular diffusion models (Kempe *et al*., 2003; Ko *et al*., 2018). With IC, a node is activated based on the active neighbouring nodes *independently* (i.e. considering the effect of each edge on the node separately). On the other hand, with LT, each node has a threshold (i.e. node weight) and it is activated if the *sum* of the weights of the edges pointing from its active neighbour nodes to this node is larger than its threshold.

To evaluate the collaboration of nodes in a network, we propose a novel algorithm to evaluate the influence of a k-seed set (i.e. k-way combination of critical nodes in our problem) on a target set (i.e. the proliferation genes) based on LT. Instead of identifying the influence spread of a k-seed set over the whole network as for solving the IM problem, we compute the influence of a k-way combination of critical nodes on a particular set of nodes in the network (i.e. all the proliferation genes in the constructed network). The detailed algorithm is illustrated in Algorithm 1 in Section 1 of the Supplement.

After applying Algorithm 1 to get the influence of k-way combinations of the top n critical nodes selected in step 3a of Stage (3) on the proliferation genes, we rank the k-way combinations in descending order of their influence. We then use the other proposed algorithm (Algorithm 2) to retain only the maximal combinations. The detail of Algorithm 2 is shown in Section 2 of the Supplement.

#### 2.2.5 Algorithms

We have developed two algorithms: Algorithm 1 for evaluating the influence of a k-seed set on a target set and Algorithm 2 for refining k-way combinations. The details of these two algorithms are in Section 1 and Section 2 of the Supplement respectively.

#### 2.2.6 Implementation

The R source code of the implementation and scripts to reproduce the experiments are available at https://github.com/pvvhoang/DriverGroup.

## 3 Results

Due to the lack of the ground truth for predicted driver gene groups, we have used several strategies to evaluate *DriverGroup*. We assess the ability of *DriverGroup* in discovering regulatory effects of gene groups in a network in Section 3.1. We evaluate the performance of *DriverGroup* in discovering driver gene groups in Section 3.2. We analyse biological implications of the predicted driver groups by using them in prognosis analysis (Section 3.3) and analysing their target genes (Section 3.4). We also use *DriverGroup* to study synthetic lethality (Section 3.5) and miRNA driver groups of EMT (Section 3.6).

### 3.1 *DriverGroup* is effective in detecting group-based regulatory effects

Before evaluating the performance of *DriverGroup* in detecting cancer driver groups, we firstly assess its performance in detecting the regulation of gene groups in a network, a key step (Step 3a) of *DriverGroup*. We use the DREAM4 data obtained from the DREAM4 In Silico Network Challenge (Marbach *et al*., 2010; Schaffter *et al*., 2011). The dataset includes 5 subsets and each subset contains the data of 100 genes, including wildtype data, knockout data (considered as expression data), dual knockout index data (i.e. indexes of 20 gene pairs which are knocked out simutaneously), dual knockout data (i.e. expression data corresponding to dual knockout index data), and network data (The detailed experiment setting is described in Section 3 of the Supplement). Given each of the 5 sub datasets, we can identify the list of genes affected by the 20 knocked out gene pairs in the network and they are considered as the gold standard of the experiment.

For each of the 5 sub datasets, we are looking at whether each method can find the targets of each knocked out pairs. We compare *DriverGroup* with jointIDA (Nandy *et al.*, 2017) and the random method. jointIDA is also used to estimate the joint effects of a group of variables on other variables. However, it estimates the joint effects of variables on a target by knocking down all variables at the same time. In the random method, we randomly pick target genes for each the knocked out gene pairs 100 times. We validate the results of each method with the gold standard above. Fig. 3 shows the precisions achieved by the three methods.

**Fig. 3.**
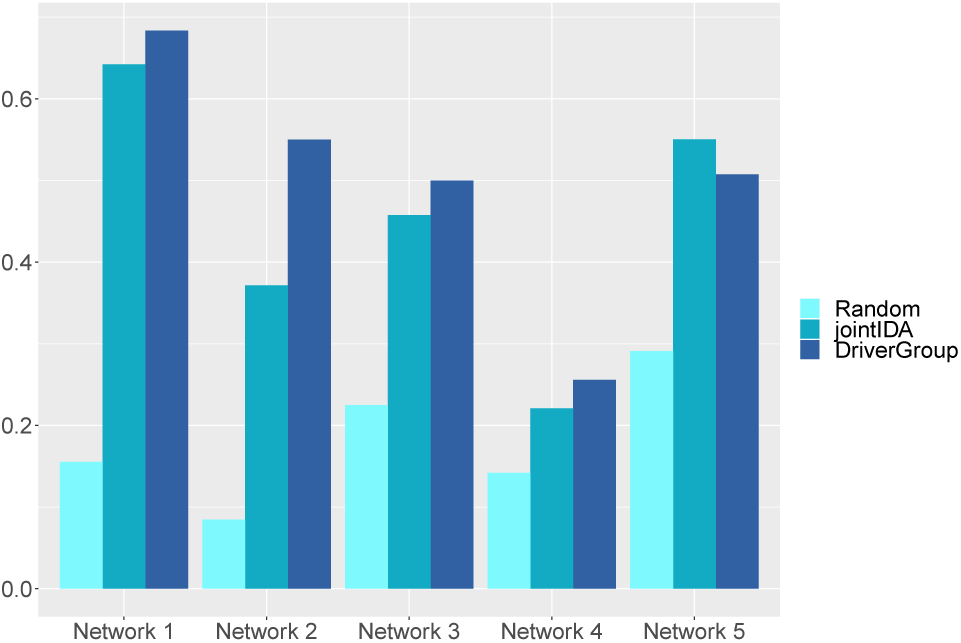
Comparison of Precision for the target genes predicted by the random method, jointIDA, and *DriverGroup* in 5 networks. The target genes predicted by each method are validated against the gold standard. Each bar indicates the *Precision* of each method.

In Fig. 3, we see that *DriverGroup* outperforms jointIDA in 4 out of the 5 cases and achieves similar precision as jointIDA in the case of network 5. Both *DriverGroup* and jointIDA outperform the random method in all cases.

To have a detailed evaluation, we compare the results of 3 methods (i.e. jointIDA, *DriverGroup*, and the random method) for all 20 gene pairs of 5 networks. For each network, we validate the predicted target genes of knocked-out gene pairs against the gold standard. We then compute the accumulated number of validated target genes of all 20 knocked-out gene pairs. The result is shown in Fig. 4. We can see that *DriverGroup* outperforms the other two methods in the first four networks and it is comparable to jointIDA in the fifth network.

**Fig. 4.**
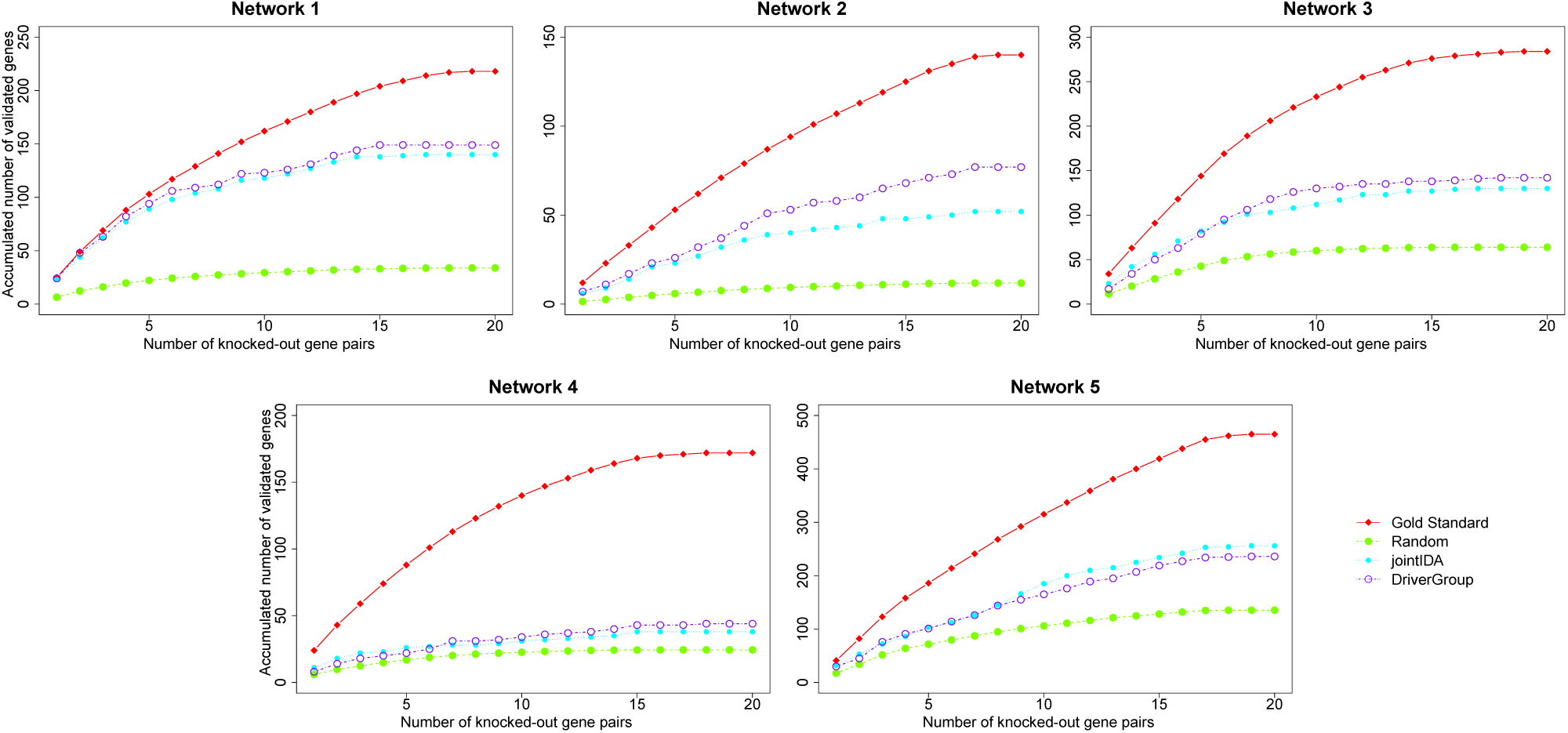
Comparison of performance of the random method, jointIDA, and *DriverGroup*. There are 5 networks in total and each chart is represented for a network. In each chart, the x-axis indicates the number of knocked-out gene pairs. The y-axis is the accumulated number of validated genes predicted by the random method, jointIDA, and DriverGroup. The red line is the gold standard and it shows the true numbers of genes affected by gene pairs. In all the cases, DriverGroup outperforms the other two methods.

In addition, the overlap of the gold standard and the target genes predicted by jointIDA and *DriverGroup* is shown in Fig. 5. In the figure, the target genes identified by jointIDA and *DriverGroup* of all 20 gene pairs of each network are validated against the gold standard. In all the 5 networks, although there are some target genes uncovered by both jointIDA and *DriverGroup*, there are a large amount of validated target genes discovered only by *DriverGroup*. Since the results of the two methods are complementary, it would be beneficial if they could be used together in predicting targets of groups of genes.

**Fig. 5.**
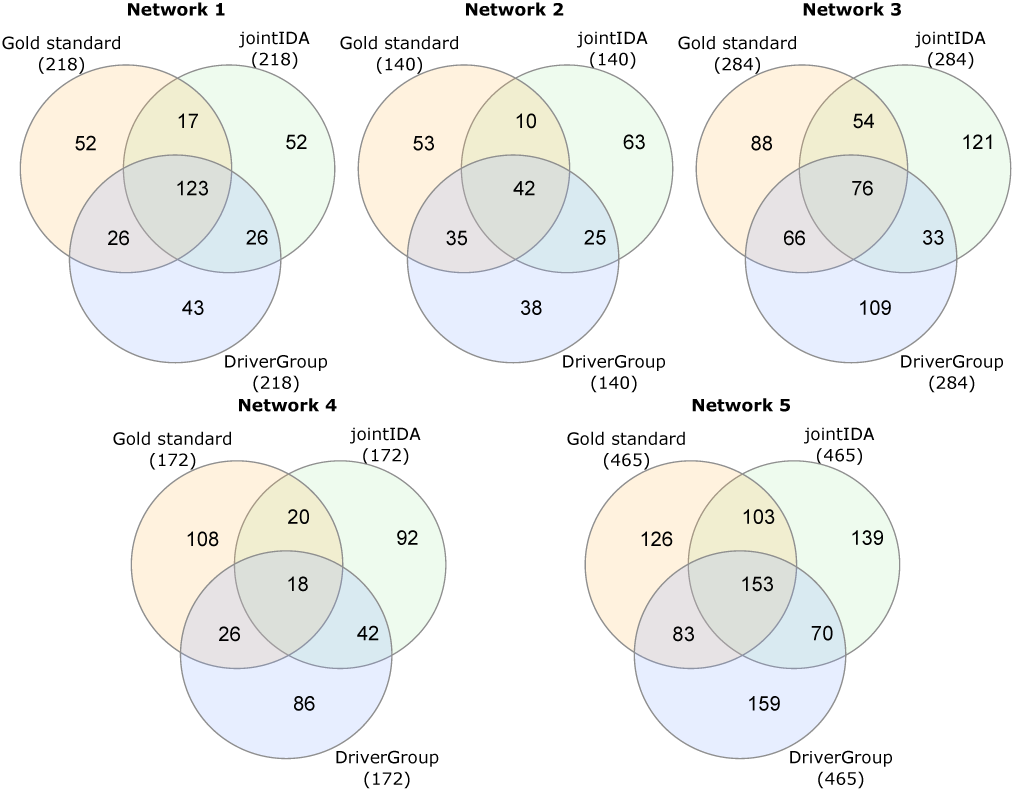
Overlap between jointIDA, *DriverGroup*, and the gold standard. The diagram shows the overlap of the gold standard and the target genes predicted by jointIDA and DriverGroup in the 5 networks. In all the five networks, DriverGroup can detect large amounts of target genes which are not discovered by jointIDA.

### 3.2 Identifying coding and miRNA cancer driver groups

We apply *DriverGroup* to the BRCA dataset to identify coding and miRNA driver groups (i.e. groups of coding RNAs/miRNAs which have an impact on the proliferation genes). We also categorise the identified groups into additional groups and enhanced groups. Additional groups regulate target genes which are in the union of the target genes of individuals in the groups. Enhanced groups regulate genes in and outside the union of the target genes of individuals in the groups. We identify 82 coding cancer driver groups and 36 miRNA cancer driver groups. We sort these groups based on their influence on the proliferation genes (i.e. The larger number of proliferation genes a group impacts on, the higher it is in the ranking list). The top 10 coding and miRNA cancer driver groups discovered by our method are presented in Table 1 and Table 2 respectively. We see that most of the identified groups are enhanced groups, indicating that members in the identified groups work collaborately to increase the effects on the proliferation genes.

**Table 1.** Coding BRCA driver groups predicted by DriverGroup. The top 10 coding driver groups are enhanced groups whose members work in concert to increase the influence on the proliferation genes.

| Group | Predicted driver groups | Size of group | Type |
| --- | --- | --- | --- |
| 1 | FOS, MBD3, JUN, E2F6, MYB, SPI1 | 6 | Enhanced |
| 2 | GATA1, FOS, MBD3, JUN, MYB, SPI1 | 6 | Enhanced |
| 3 | TCF3, FOS, MBD3, JUN, MYB, SPI1 | 6 | Enhanced |
| 4 | GATA1, FOS, MBD3, JUN, SPI1 | 5 | Enhanced |
| 5 | GATA1, TCF3, FOS, MBD3, JUN, MYB | 6 | Enhanced |
| 6 | GATA1, FOS, MBD3, SPI1 | 4 | Enhanced |
| 7 | FOS, MBD3, JUN, SPI1 | 4 | Enhanced |
| 8 | TCF3, FOS, MBD3, JUN, E2F6, MYB | 6 | Enhanced |
| 9 | GATA1, MBD3, JUN, MYB | 4 | Enhanced |
| 10 | MBD3, JUN, MYB, SPI1 | 4 | Enhanced |

**Table 2.** miRNA BRCA driver groups predicted by DriverGroup. The top 10 miRNA driver groups are enhanced groups whose members work in concert to increase the influence on the proliferation genes.

| Group | Predicted driver groups | Size of group | Type |
| --- | --- | --- | --- |
| 1 | hsa-miR-96-5p, hsa-miR-342-5p | 2 | Enhanced |
| 2 | hsa-miR-130a-5p, hsa-miR-34a-5p, hsa-miR-22-5p, hsa-miR-222-5p, hsa-miR-223-5p | 5 | Enhanced |
| 3 | hsa-miR-130a-5p, hsa-miR-22-5p, hsa-miR-222-5p, hsa-miR-223-5p, hsa-miR-6797-5p | 5 | Enhanced |
| 4 | hsa-miR-130a-5p, hsa-miR-34a-5p, hsa-miR-22-5p | 3 | Enhanced |
| 5 | hsa-miR-130a-5p, hsa-miR-22-5p, hsa-miR-146a-5p | 3 | Enhanced |
| 6 | hsa-miR-130a-5p, hsa-miR-22-5p, hsa-miR-6797-5p | 3 | Enhanced |
| 7 | hsa-miR-34a-5p, hsa-miR-22-5p, hsa-miR-222-5p | 3 | Enhanced |
| 8 | hsa-miR-34a-5p, hsa-miR-22-5p, hsa-miR-223-5p | 3 | Enhanced |
| 9 | hsa-miR-34a-5p, hsa-miR-22-5p, hsa-miR-6797-5p | 3 | Enhanced |
| 10 | hsa-miR-130a-5p, hsa-miR-34a-5p, hsa-miR-222-5p, hsa-miR-342-5p, hsa-miR-6797-5p | 5 | Enhanced |

The driver groups predicted by *DriverGroup* are promising as some members of the predicted groups are confirmed to be related to breast cancer. Among the genes in the top 10 coding cancer driver groups predicted by *DriverGroup*, GATA1, TCF3, JUN, and MYB are in the Cancer Gene Census (CGC) from the COSMIC database (Forbes *et al*., 2015). In addition, other genes, including FOS, MBD3, E2F6, and SPI1, are also previously proved to be related to breast cancer. Specifically, FOS is critical to the growth of MCF-7 breast cancer cell (Lu *et al*., 2005) and its family plays an important role in the biological function of breast tumours (Langer *et al*., 2006). There is a relationship between MBD3 and human breast cancer cells (Shimbo *et al*., 2016). E2F6 regulates BRCA1 negatively in human cancer cells (Oberley *et al*., 2003) and SPI1 can be used for prognosis in breast cancer (Wang *et al*., 2007).

Among the miRNAs in the top 10 miRNA cancer driver groups predicted by *DriverGroup*, there are 3 miRNAs (hsa-miR-22-5p, hsa-miR- 342-5p, and hsa-miR-34a-5p) involved in tumorigenesis of breast cancer, which are confirmed by OncomiR (Wong *et al*., 2018), a database for studying pan-cancer miRNA dysregulation. Out of these 3 miRNAs, hsamiR- 342-5p is proved to be a regulator of the development of breast cancer cells in another work (Lindholm *et al*., 2018) as well. Another 3 miRNAs, hsa-miR-130a-5p, hsa-miR-146a-5p, and hsa-miR-223-5p, are also confirmed to be related to breast cancer. Specifically, hsa-miR-130a-5p targets FOSL and upregulates ZO-1 to suppress breast cancer cell migration (Chen *et al*., 2018), hsa-miR-146a-5p has an over expression in breast cancer cells (Sandhu *et al*., 2014), and hsa-miR-223-5p is a coordinator of breast cancer (Pinatel *et al*., 2014).

### 3.3 Predicted driver groups are useful in predicting survival

As the predicted driver groups likely cause carcinogenesis, they could be promising biomarkers for tumour classification. To explore this concept, we use the driver gene groups predicted by *DriverGroup* to stratify breast cancer patients. We obtain the BRCA gene expression data from Zhang *et al.* (2019), which includes clinical data, for survival analysis. We use the first predicted coding driver groups in Table 1, including FOS, MBD3, JUN, E2F6, MYB, and SPI1, and the Similarity Network Fusion (SNF) method (Xu *et al.*, 2017; Wang *et al.*, 2014) to cluster cancer patients (see Section 4 of the Supplement for the results with the second and the third driver groups). SNF takes expression of these genes (i.e. 6 genes in this case) as input and outputs subtypes of cancer patients. We then evaluate the survival outcomes of patients in the classified subtypes. The results show that the survivals of patients in different subtypes are significantly different (p-value = 0.0152) as in Fig. 6. In addition, the clustering display indicates the similarity of samples in each subtype and the silhouette plot shows a high quality clustering with a large average silhouette width (i.e. 0.77).

**Fig. 6.**
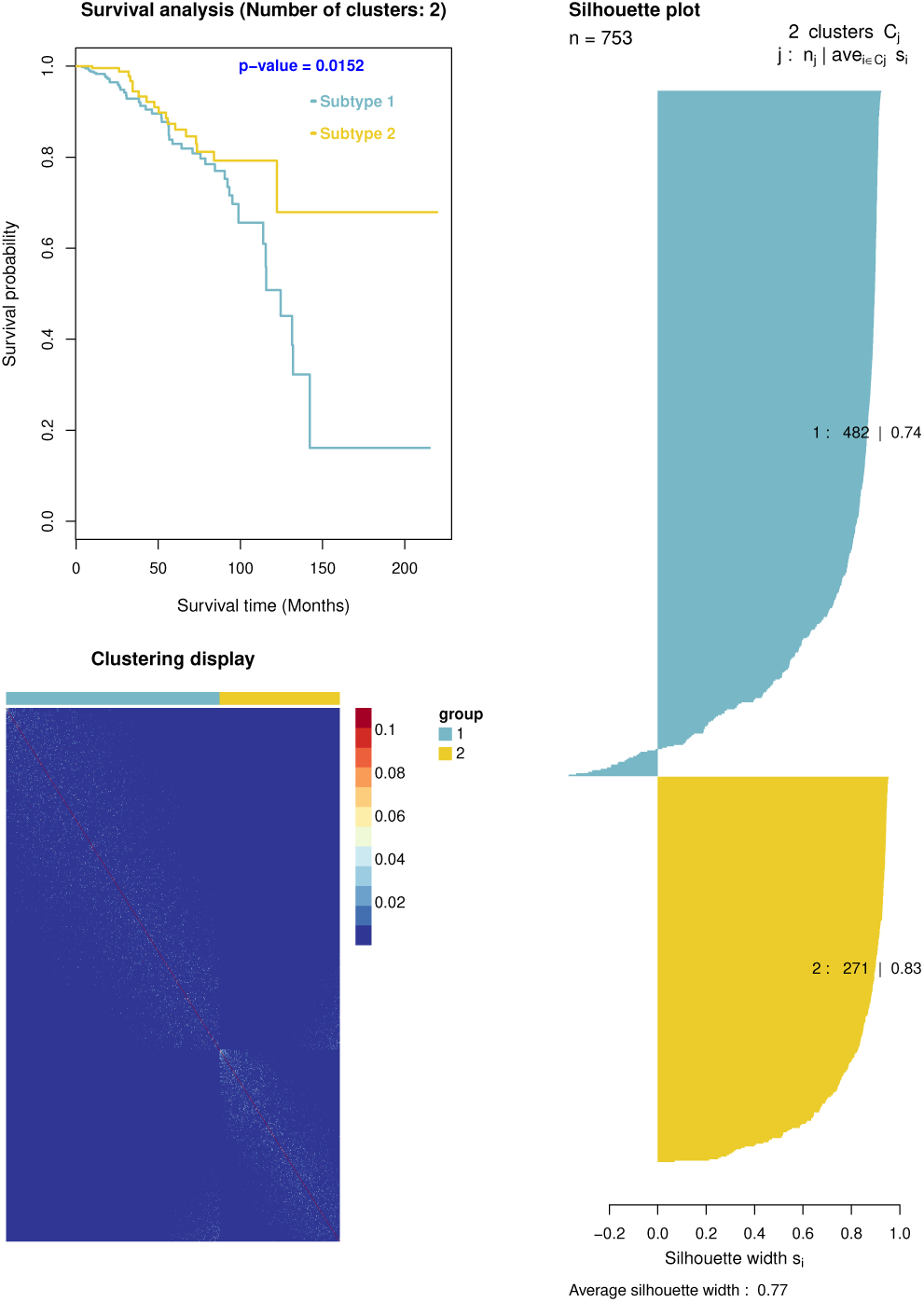
Survival curves, silhouette plot, and clustering display. Survival curves, silhouette plot, and clustering display of cancer subtypes identified by using *the first predicted coding driver groups (including FOS, MBD3, JUN, E2F6, MYB, and SPI1)* indicate that the survivals of patients are significantly different in the two subtypes and the clustering is highly qualified with a large average silhoutte width and the similarity of samples in each subtype.

### 3.4 Members of predicted driver groups regulating common target genes

To see the functional association among the members of driver groups predicted by *DriverGroup*, we check if they regulate common target genes. We use the TransmiR database of TF-miRNA interactions to identify target genes of the members of predicted coding driver groups and use the miRTarBase, TarBase, miRWalk, and TargetScan databases of miRNA-TF/mRNA interactions to identify target genes of predicted miRNA driver groups. We observe that for both the top 10 predicted coding and noncoding driver groups, all participants in each group regulate some common target genes, indicating the functional link of the members in driver groups identified by our proposed method.

### 3.5 Detecting synthetic lethality with *DriverGroup*

Two genes have a synthetic lethal (SL) interaction if the perturbation of both genes simultaneously is lethal but a perturbation that affects either gene alone is viable (Lord *et al.*, 2015; O’Neil *et al.*, 2017). It means that in cancer patients, the collaboration of two genes in a SL interaction results in the loss of viability. Applying *DriverGroup*, we identify SL interactions by detecting only the driver gene groups of size 2. We validate the top 1,000 predicted SL gene pairs against SynLethDB (Guo *et al.*, 2016), an existing synthetic lethality database. There are 6 validated SL gene pairs. Based on the hypergeometric test, the overlap between the predicted SL gene pairs and the gold standard is significant, with a p-value of 0.00027.

### 3.6 Detecting driver groups of EMT

Metastasis is a process where cancer cells migrate from the primary tumour to distant locations in the body. It is the major cause of death of cancer patients. EMT is one of the processes which create these metastatic cells (Park *et al.*, 2008). EMT is promoted by coding genes (Lee *et al.*, 2018) and/or non-coding genes (Gregory *et al.*, 2008). In this section, we apply *DriverGroup* to the BRCA dataset to discover driver groups for the EMT of breast cancer patients by identifying miRNA groups which have maximum influence on the EMT signatures (Tan *et al.*, 2014). As *DriverGroup* detects miRNA groups which regulate EMT signatures, the detected miRNA groups are expected to drive the EMT transition in breast cancer patients. We identify 61 miRNA driver groups for EMT and we sort these groups based on their influence on the EMT signatures (i.e. The larger number of EMT signatures a group impacts on, the higher it is in the ranking list). The list of top 10 miRNA driver groups for EMT in breast cancer is shown in Table 3. Among these miRNAs, hsa-miR-130a-5p and hsa-miR-223-5p are EMT miRNAs (Cursons *et al.*, 2018), indicating the potential of *DriverGroup* in detecting driver groups for different biological processes such as EMT.

**Table 3.**
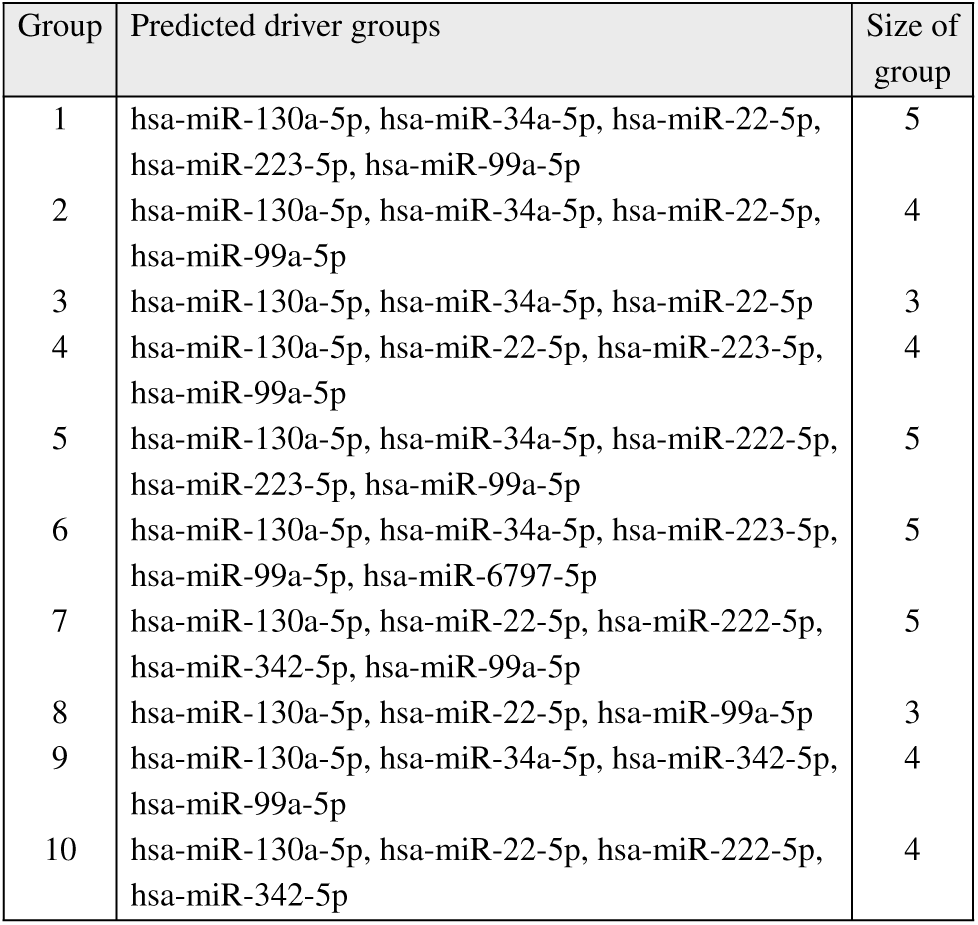
miRNA driver groups of EMT predicted by DriverGroup

| Group | Predicted driver groups | Size of group |
| --- | --- | --- |
| 1 | hsa-miR-130a-5p, hsa-miR-34a-5p, hsa-miR-22-5p, hsa-miR-223-5p, hsa-miR-99a-5p | 5 |
| 2 | hsa-miR-130a-5p, hsa-miR-34a-5p, hsa-miR-22-5p, hsa-miR-99a-5p | 4 |
| 3 | hsa-miR-130a-5p, hsa-miR-34a-5p, hsa-miR-22-5p | 3 |
| 4 | hsa-miR-130a-5p, hsa-miR-22-5p, hsa-miR-223-5p, hsa-miR-99a-5p | 4 |
| 5 | hsa-miR-130a-5p, hsa-miR-34a-5p, hsa-miR-222-5p, hsa-miR-223-5p, hsa-miR-99a-5p | 5 |
| 6 | hsa-miR-130a-5p, hsa-miR-34a-5p, hsa-miR-223-5p, hsa-miR-99a-5p, hsa-miR-6797-5p | 5 |
| 7 | hsa-miR-130a-5p, hsa-miR-22-5p, hsa-miR-222-5p, hsa-miR-342-5p, hsa-miR-99a-5p | 5 |
| 8 | hsa-miR-130a-5p, hsa-miR-22-5p, hsa-miR-99a-5p | 3 |
| 9 | hsa-miR-130a-5p, hsa-miR-34a-5p, hsa-miR-342-5p, hsa-miR-99a-5p | 4 |
| 10 | hsa-miR-130a-5p, hsa-miR-22-5p, hsa-miR-222-5p, hsa-miR-342-5p | 4 |

## 4 Conclusion

Since there is evidence showing that genes work in concert to regulate targets and progress cancer, several methods have been developed to identify these genes. However, current methods only discover mutated modules. Only one mutated gene in a module is sufficient to progress cancer. Thus, members in these mutated modules do not collaborate in driving cancer and these mutated modules are not truly driver gene groups. In addition, current methods only identify coding drivers while non-coding genes can also regulate targets to drive cancer. Therefore, novel methods are required to identify driver gene groups to elucidate their regulatory mechanism.

To overcome the limitations of existing methods, in this paper, we have developed a novel method, *DriverGroup*, to uncover coding and non-coding cancer driver groups. We have evaluated the effectiveness of *DriverGroup* with various experiments. The results have demonstrated that *DriverGroup* can explore promising driver gene groups. Predicted coding driver groups can be used to classify cancer patitents into subtypes and the survivals of patients in different subtypes are significantly different. Furthermore, *DriverGroup* can also detect synthetic lethal gene pairs and EMT driver groups. All these results show that the findings of *DriverGroup* can provide new insights into molecular regulatory mechanisms of cancer initialisation and progression, and *DriverGroup* has the potential to contribute to the development of effective cancer treatments.

## Acknowledgements

This research is supported by the Australian Government Research Training Program (RTP) Scholarship and the Vice Chancellor & President’s Scholarship offered by the University of South Australia.

